# Nasal BCG Exposure Accelerates Dural Lymphatics Development via Macrophages’ Role in Newborn Mice

**DOI:** 10.1101/2021.04.27.441593

**Authors:** Junhua Yang, Lifang Yuan, Linyang Song, Fangfang Qi, Zejie Zuo, Jie Xu, Zhibin Yao

## Abstract

The dural lymphatics develop mainly during the first postnatal month. Lymphatics may be shaped by immune activation when bacterial infection happens. BCG, a strong immune activator, is widely injected to newborns. Moreover, human beings are nasally exposed in daily life to bacterial stimuli. These background prompted us to investigate whether neonatal BCG injection combined with nasally exposure exerts an influence on dural lymphatics develop. Here, mice received a single dose intracutaneous (i.c.) BCG injection immediately after birth followed by repeated nasal BCG challenge once a day (i.c./nasal group). These mice had an accelerated dural lymphatics growth and increased levels of several cytokines and VEGFR-3. Furthermore, macrophages were identified as a key mediator of these alterations. Mice that received mere one dose i.c. BCG injection showed no significant alterations in these indexes. Additionally, mere repeated nasal BCG challenge induced similar effects to i.c./nasal challenge but with a slighter extent. Taken together, these findings show that repeated nasal BCG vaccination accelerates dural lymphatics development in neonatal mice, especially in the presence of neonatal i.c. BCG injection.

## Introduction

Since two publications in 2015 describing the structure and function of the dural lymphatics (*1, 2*), these vessels have drawn increasing attention. A series of researches as well as review articles focused on them (*3, 4*). In 2017, Salli Antila et al. reported the development and plasticity of dural lymphatics in mice, revealing that the dural lymphatics network starts to develop after birth and they were already well developed at about four weeks old (*5*). As well known, lymphatics may be shaped by inflammatory factors produced in immune activation. It has been reported that immune activation induced by bacterial or virus infection led to lymphangiogenesis outside the central nervous system (*6–8*). BCG is a live attenuated vaccine containing live *Mycobacterium tuberculosis* and is widely used to newborns. Planned BCG vaccination is a strong and unnatural neonatal immune stimuli. About 90 percent of infants of the world were administrated neonatally with BCG (*9*).

We here imitated the immune challenge by BCG that happens to human infants, by means of a single dose i.c. BCG injection immediately after birth. Moreover, we also imitated the daily nasal exposure to microbiota that exist in the air by repeated nasal challenge with BCG performed once a day within the first 14, 19 or 21 postnatal days. For the nasal immune challenge with bacteria, BCG was used for three reasons: 1) this constitutes a boosting immunization when combined with neonatal i.c. BCG injection in spite of a different exposure pathway, 2) in the long period of human evolution and survival, *Mycobacterium tuberculosis*, as a kind of bacteria mainly transmitted through the air, has been exposed to almost all human individuals (*10*), 3) despite advances in modern medicine, one third of the world’s population is estimated to have contracted the bacteria, with estimates of ten million new infections globally each year (*11*). Meanwhile, mice receiving PBS, mere one dose i.c. neonatal BCG injection or mere repeated nasal BCG challenge were also prepared and observed. We found that neonatal BCG vaccination accelerates lymphatics development in mice, especially in the presence of neonatal i.c. BCG exposure.

## Materials and methods

### Animals and study design

Newborn litters of C57Bl/6 mice were obtained from the SYSU Laboratory Animal Center (Guangzhou, China) and were housed in a specific pathogen-free facility. This study consisted of different tests, each of which used four groups of mice, namely con-group, intracutaneous(i.c.)-group, nasal-group and i.c./nasal-group. All experiments in the study were started on postnatal day 0 (P0).

For each test, six newborn litters of C57Bl/6 mice were used. Within each litter, four male pups were used and randomly divided into four groups, receiving i.c. PBS injection combined with nasal PBS administration (con-group), i.c. BCG injection combined with nasal PBS administration (i.c.-group), i.c. PBS injection combined with nasal BCG vaccination (nasal-group) or i.c. BCG injection combined with nasal BCG vaccination (i.c./nasal-group). The pups that were sacrificed after P21 were weaned on P21. The study was approved by the SYSU Institute Research Ethics Committee and all animal experiments were conducted in compliance with the relevant laws and institutional guidelines.

### Immunization procedures

A single dose of BCG was administered (i.c.) to the mice in i.c.-group and i.c./nasal-group in the back on P0. For nasal-group and i.c./nasal-group, each mouse received nasal administration of BCG once a day from P0 to P14, P19 or P21. Freeze-dried living BCG (D2-BP302 strain, Biological Institute of Shanghai, China) was suspended in PBS. The dose both of i.c. injection and nasal administration is about 10^5^ CFU.

### Immunofluorescence staining and quantification

To test the proliferating cells in dural lymphatics, the mice received five intraperitoneal injections of BrdU (Sigma-Aldrich, 50 mg/kg) from P14 once a day to label the dividing cells. At P19, the animals were deeply anesthetized with chloral hydrate (10% solution) and then transcardially perfused with PBS. After removing the mandibles and the skull rostral to the maxillae, the skull cap and the attached dura/arachnoid containing superior sagittal sinus (SSS) and two transverse sinus (TS) areas were removed with surgical scissors. The dura mater tissue used for all the other tests were obtained from mice at P14 or P21 that were not treated with BrdU. Whole mounted meninges that were still attached to the skull cap were placed in 4% paraformaldehyde (PFA) in PBS and postfixed for 24 h at 4°C. The dura/arachnoid were then carefully and gently separated from the skull cap using tweezers with a thin head. Actually, the adhesion between dura mater and calvaria is relatively loose in young mice, and therefore it is easy to keep them intact during peeling from calvaria. The dura were then washed with PBS three times and processed for staining. The dura were blocked in PBS containing 1% bovine serum albumin and 0.25% Triton X-100 (Sigma-Aldrich, St. Louis, MO, USA) for 1 h at 37°C before being incubated with primary antibodies diluted in the same blocking solution overnight at 4°C. The primary antibodies were as follows: goat anti-Lyve-1 (R&D Systems, Minneapolis, MN, USA; 1:600), rabbit anti-Prox-1 (Abcam, Cambridge, MA, USA; 1:500), rat anti-BrdU (Santa Cruz, CA, USA;1:500), goat anti-VEGFR-3 (R&D Systems, Minneapolis, MN, USA; 1:50) and rat anti-F4/80 (Abcam, Cambridge, MA, USA; 1:200). Specimens were washed three times with PBS before being incubated with the following secondary antibodies for 2 h at 37°C: Alexa Fluor 555-conjugated donkey anti-goat, Alexa Fluor 647-conjugated donkey anti-rat and Alexa Fluor 488-conjugated donkey anti-rabbit antibodies. All secondary antibodies (Invitrogen Molecular Probes^®^, Eugene, Oreg., USA) were diluted 1:400. Each of the dura samples was then placed on a glass slide with the skull face attached on the slide. This process was performed in PBS to ensure that the SSS area and two TS areas stretched without folding.

Confocal micrographs of immunofluorescence staining were obtained using a LSM 780 confocal laser-scanning microscope (Zeiss, Heidelberg, Germany). The total numbers of each kind of labeled cells showed in Results section within the selected areas were quantified using a Stereo Investigator (MicroBrightField, Williston, USA) stereology system by an investigator who was blinded to the groups.

### Multiplex assay

Mice at P21 were anesthetized with chloral hydrate (10% solution). Then, 5-7 μl of cerebrospinal fluid (CSF) were obtained from the cisterna magna of each mouse, according to a previously described protocol (*12*). The mouse cytokine/chemokine panel (MCYTOMAG-70K-04; Millipore, Billerica, MA, USA) and bead-based Luminex system were used to detect the concentrations of IFN-γ, IL-4, TNF-α, IL-7, IL-17, IL-1β, IL-6, IL-10 and monocyte chemotactic protein-1 (MCP-1) in the CSF. We first added 25 μl of prepared CSF samples that had previously been diluted 1:5 in assay buffer to each well. Then, 25 μl of the magnetic beads coated with specific antibodies were added, and the mixture was incubated overnight at 4°C. Then, the beads were washed and incubated with 25 μl of the biotinylated detection antibody for 1 h at room temperature (RT). Afterwards, 25 μl of the streptavidin-phycoerythrin conjugate were added, and the reaction was incubated for 30 min at RT. The beads were then washed with 150 μl of sheath fluid for 5 min at RT. Data were collected on a Bio-Plex-200 system (Bio-Rad, Hercules, CA, USA) and analyzed using professional software (Bio-Plex Manager; Bio-Rad, CA, USA).

### ELISA

Mice at P21 were anesthetized with chloral hydrate (10% solution). Then, 5-7 μl of CSF were obtained from the cisterna magna of each mouse, according to a previously described protocol (*12*). VEGF-C levels were assayed in the CSF of mice using an ELISA kit (VEGF-C, Cusabio, China), according to the manufacturer’s instruction.

### Western blot

At P21, levels of the VEGFR-3 proteins in the dura were examined by western blot analyses. After the animals were anesthetized and perfused with PBS, the skull cap and the attached dura/arachnoid were obtained. Before and after the dura/arachnoid were separated from the skull cap, the tissues were washed three times with ice-cold PBS to remove the CSF and intercellular fluid that may have adhered to the meninges. Then, the dura/arachnoid were homogenized in ice-cold RIPA buffer. The homogenate was centrifuged at 9,000 g for 15 minutes at 4°C, after which the supernatant was analyzed by western blotting. Protein concentrations were quantified using the bicinchoninic acid (BCA) protein assay kit (Biotime, China). Membrane homogenates (50 μg) were separated on 6% SDS-PAGE gels and transferred onto presoaked PVDF membranes (Bio-Rad, Pleasanton, CA, USA). Blots were then blocked with 10% non-fat milk in PBST (100 nM phosphate buffer, pH 7.5, containing 150 nM NaCl and 0.1% Tween-20) for 2 h at RT. The the membranes were incubated overnight at 4°C with rabbit anti-VEGFR-3 antibodies (Abcam, Cambridge, MA, USA; 1:50). Horseradish peroxidase (HRP)-conjugated secondary goat anti-rabbit antibodies (Sigma-Aldrich Co., USA) were used at a dilution of 1:4000. Protein bands were visualized using a chemiluminescence detection system (Pierce, Milan, Italy). Optical densities were measured by dividing the intensity of the bands by the intensity of the housekeeping protein β-actin (Cell Signaling Technology, MA, USA, 1:1000). Data from control samples were assigned a relative value of 100%.

### Statistical analyses

All data were statistically analyzed using the Statistical Package for the Social Sciences 25.0. The data were analyzed using randomly completely block design ANOVA followed by Bonferroni’s *post hoc* test or paired-samples *t* test. The statistically significant level was set at *α* = 0.05. The normality and homogeneity of variance were conducted using the Shapiro-Wilk test and Levene’s test, respectively.

## Results

### Accelerated dural lymphatics development in nasal-group and i.c./nasal-group in TS zone at P14

At P14, lymphatics distribution around the TS was observed by immunofluorescencely staining for Lyve-1. Lyve-1^+^ vessels covered about 59%, 63%, 85% and 100% of the length of ipsilateral TS in con-group, i.c.-group, nasal-group and i.c./nasal-group, respectively (Fig.1). Compared with con-group, both nasal-group and i.c./nasal-group had a significant increase in the length of TS covered by Lyve-1^+^ vessels, Lyve-1^+^ vessels area, numbers of sprouting from Lyve-1^+^ vessels and numbers of cross points of Lyve-1^+^ vessels (Fig.1). In all these four indexes, the i.c./nasal-group possessed a significant larger extent of increase than the nasal-group (Fig.1). Moreover, the i.c./nasal-group showed obvious sprouting and extending of lymphatics from the TS zone toward both rostral (upward) and caudal side (downward) in the dura mater (Fig.1D). All these findings indicated accelerated dural lymphatics development by neonatal nasal exposure to BCG.

**Fig.1.**
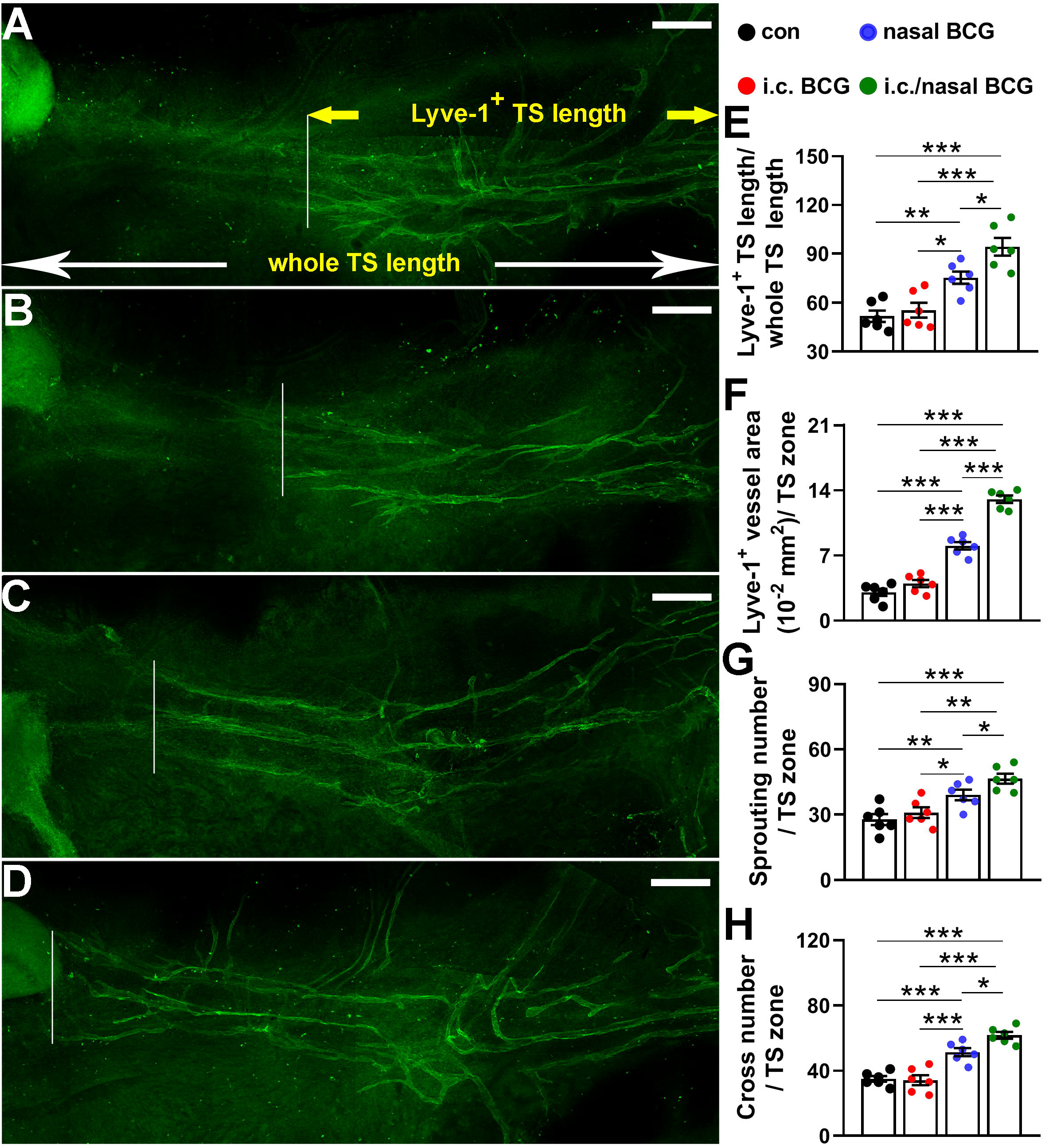
Accelerated dural lymphatics development in nasal-group and i.c./nasal-group mice at P14. (A-D) Representative graphs show the differences in dLVs within the corresponding TS areas of the dura in each group of mice. Scale bar, 200 μm. (E-H) Bars represent the length of TS covered by Lyve-1^+^ vessels, Lyve-1^+^ vessels area%, numbers of sprouting from Lyve-1^+^ vessels and numbers of cross points of Lyve-1^+^ vessels dLVs within the unilateral TS area. *n* = 6 mice/group. *: *p* < 0.05; **: *p* < 0.01; ***: *p* < 0.001. Randomly completely block design ANOVA followed by Bonferroni’s *post hoc* test. Data are presented as the mean ± SEM.

### Accelerated dural lymphatics development with more macrophage-derived lymphatic endothelial cell progenitors (M-LECPs) in SSS zone in nasal-group and i.c./nasal-group at P21

Macrophages (F4/80^+^) play a vital role in lymphangiogenesis induced by immune stimuli because of their capacity to transform into lymphatic endothelial progenitors (*13, 14*). F4/80^+^/Lyve-1^+^ cells, F4/80^+^/Prox-1^+^ cells and F4/80^+^/Lyve-1^+^/Prox-1^+^ cells have previously been reported as the macrophage-derived LEC progenitors (*13, 14*).

In the current study, lymphatics distribution and macrophages presence around the SSS was observed after the dura mater was immunofluorescencely trible-stained for Lyve-1/Prox-1/F4/80 at P21. Lyve-1^+^ vessels covered about 30%, 50%, 70% and 100% of the length of SSS in con-group, i.c.-group, nasal-group and i.c./nasal-group, respectively (Fig.2). Compared with con-group, both nasal-group and i.c./nasal-group had a significant increase in Lyve-1^+^ vessels area%, numbers of sprouting from Lyve-1^+^ vessels and Prox-1^+^ nucleus numbers (Fig.2). In all these three indexes, the i.c./nasal-group possessed a significant larger extent of increase than the nasal-group (Fig.2). These indexes also indicated accelerated dural lymphatics development by BCG, especially considering the fact that lymphatics begin to cover 100% of the length of SSS at P28 under normal condition (*5*).

**Fig.2.**
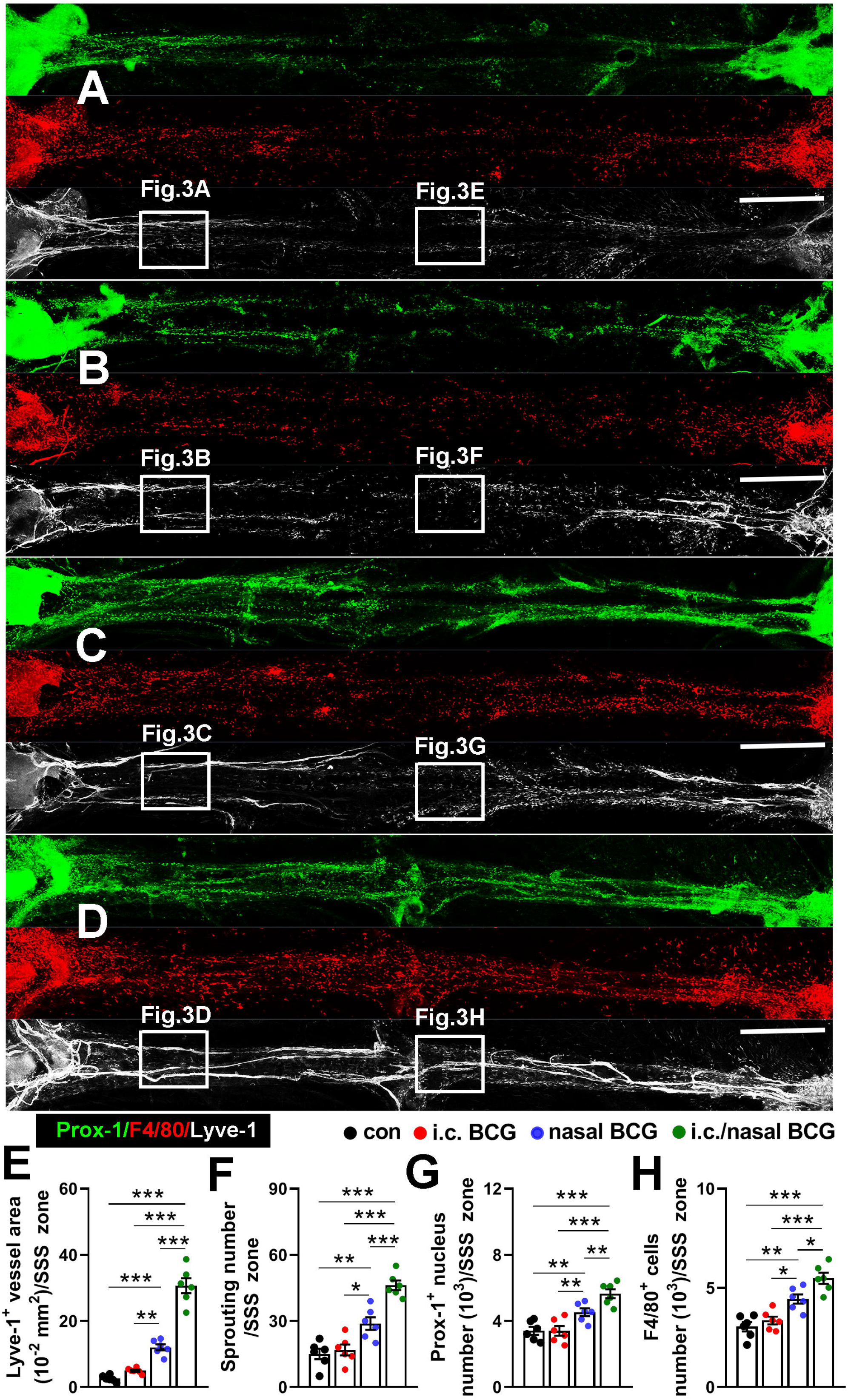
Accelerated dural lymphatics development with more M-LECPs in SSS zone in nasal-group and i.c./nasal-group at P21. (A-D) Representative graphs show the differences in dLVs distribution and macrophages number within the corresponding SSS areas of the dura in each group of mice. Scale bar, 400 μm. (E-H) Bars represent Lyve-1^+^ vessels area, numbers of sprouting from Lyve-1^+^ vessels, Prox-1^+^ nucleus numbers and the number of total F4/80^+^ cells. *n* = 6 mice/group. *: *p* < 0.05; **: *p* < 0.01; ***: *p* < 0.001. Randomly completely block design ANOVA followed by Bonferroni’s *post hoc* test. Data are presented as the mean ± SEM.

Compared with con-group, both nasal-group and i.c./nasal-group had a significant increase in the number of total F4/80^+^ cells (Fig.2H), F4/80^+^/Prox-1^+^ cells and F4/80^+^/Prox-1^+^/Lyve-1^+^ cells (Fig.3M-O). In all these three indexes, the i.c./nasal-group possessed a significant larger extent of increase than the nasal-group (Fig.2H) (Fig.3M-O). In addition, M-LECPs in different stages were found, indicating by different features of phenotype and location to lymphatics (See detailed description in Fig.4 and the corresponding legend).

**Fig.3.**
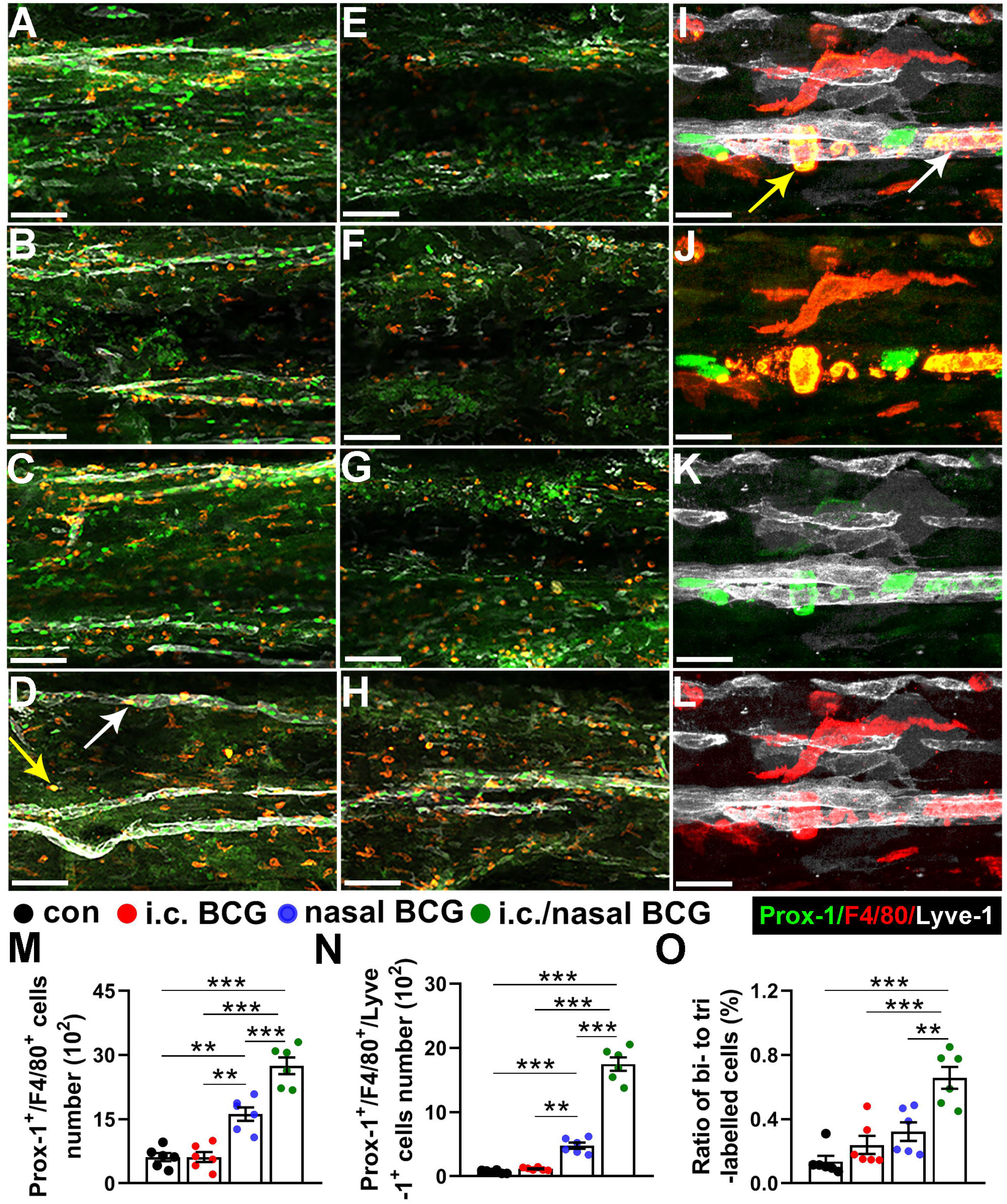
Selected representative scopes from graphs in Fig.2A-D, showing dural lymphatics and M-LECPs presence in SSS zone in nasal-group and i.c./nasal-group at P21. (A-D) Selected representative scopes that were next to the sinus confluence (the left end) of each SSS graph in Fig.2A-D. Scale bar, 100 μm. (E-H) Selected representative scopes that were located at the middle of each SSS graph in Fig.2A-D. Scale bar, 100 μm. (I-L) Representative high-magnified scopes showing co-labelled Lyve-1^+^ (white), Prox-1^+^(green) and F4/80^+^ (red) signals. (M-N) Bars represent the numbers of F4/80^+^/Prox-1^+^ cells and F4/80^+^/Prox-1^+^/Lyve-1^+^ cells within the whole SSS area of each group. (O) The ratio of F4/80^+^/Prox-1^+^/Lyve-1^+^ cells numbers to F4/80^+^/Prox-1^+^ cells numbers within the whole SSS area of each group. *n* = 6 mice/group. *: *p* < 0.05; **: *p* < 0.01; ***: *p* < 0.001. Randomly completely block design ANOVA followed by Bonferroni’s *post hoc* test. Data are presented as the mean ± SEM.

**Fig.4.**
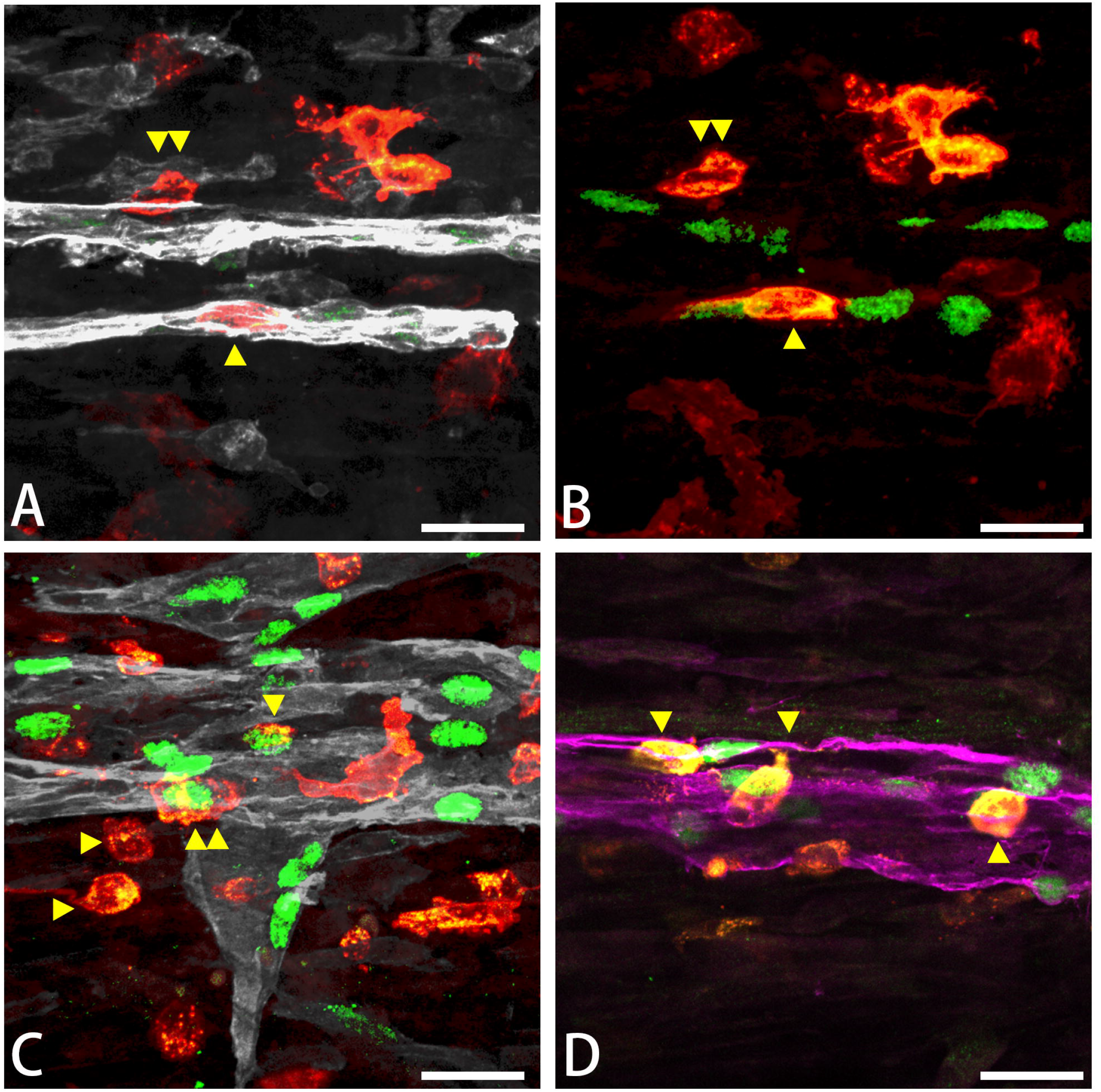
Selected representative scopes showing the various stages of M-LECPs in their transformation into dural lymphatics in SSS zone. (A-B) The same representative scope showing co-labelled Lyve-1^+^ (white), Prox-1^+^(green) and F4/80^+^ (red) signals. (C) Another representative scope showing co-labelled Lyve-1^+^ (white), Prox-1^+^(green) and F4/80^+^ (red) signals. (D) Another representative scope showing co-labelled Lyve-1^+^ (purple), Prox-1^+^(green) and F4/80^+^ (red) signals.

### Elevated real time lymphangiogenesis level in SSS zone in nasal-group and i.c./nasal-group at P19

The real time lymphangiogenesis level was also evaluated by immunofluorescencely co-staining for Lyve-1/Prox-1/BrdU at P19 and the SSS zone of dura mater was observed. Fig.S1A-D shows whole length graph of SSS area from which representative higher scopes were presented in Fig.5. As expected, both nasal-group and i.c./nasal-group had a significant increase in BrdU^+^ cells (Fig. S1F), BrdU^+^/Prox-1^+^ (Fig.5M) and BrdU ^+^/Prox-1^+^/Lyve-1^+^ (Fig.5N) cells within the entire area of SSS compared with con-group besides the increased indexes that reflecting the accelerated dural lymphatics development (Fig. S1E, G-H). In both indexes, the i.c./nasal-group possessed a significant larger extent of increase than the nasal-group.

**Fig.5.**
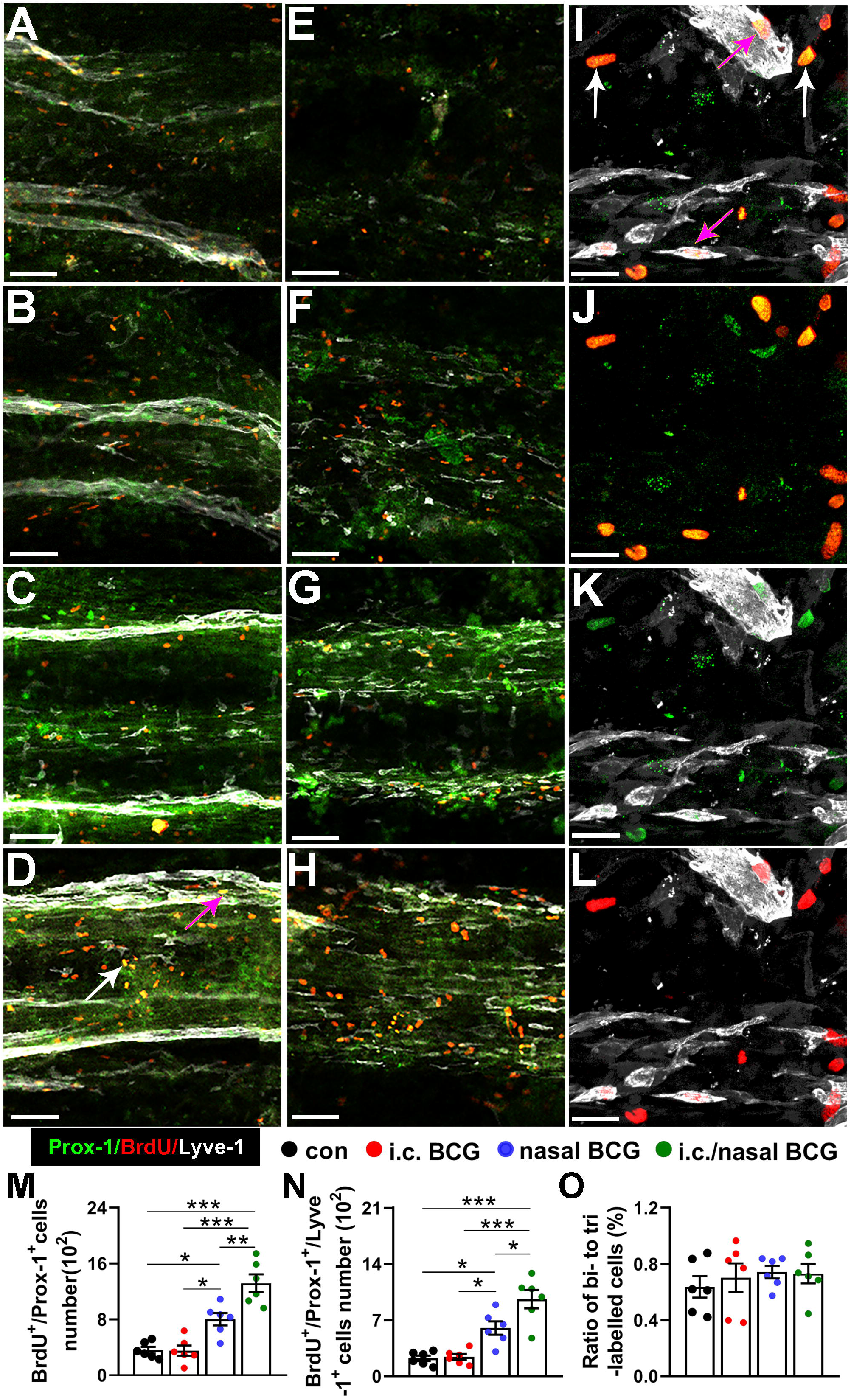
Selected representative scopes from graphs in Fig. S1A-D, showing the elevated real time lymphangiogenesis level in SSS zone in nasal-group and i.c./nasal-group. (A-D) Selected representative scopes that were next to the sinus confluence of each SSS graph in Fig. S1A-D. Scale bar, 50 μm. (E-H) Selected representative scopes that were located at the middle of each SSS graph in Fig. S1A-D. Scale bar, 50 μm. (I-L) Representative high-magnified scopes showing co-labelled Lyve-1^+^ (white), Prox-1^+^(green) and BrdU^+^ (red) signals. (M-N) Bars represent the numbers of BrdU^+^/Prox-1^+^ cells and BrdU^+^/Prox-1^+^/Lyve-1^+^ cells within the whole SSS area of each group. *n* = 6 mice/group. (O) The ratio of BrdU^+^/Prox-1^+^/Lyve-1^+^ cells numbers to BrdU^+^/Prox-1^+^ cells numbers within the whole SSS area of each group. *: *p* < 0.05; **: *p* < 0.01; ***: *p* < 0.001. Randomly completely block design ANOVA followed by Bonferroni’s *post hoc* test. Data are presented as the mean ± SEM.

### Pro-lymphangiogenesis and macrophage-recruting profile of cytokine expression in the CSF from mice in nasal-group and i.c./nasal-group at P21

Lymphangiogenesis is a dynamic process that is regulated also by other cytokines, such as vascular endothelial growth factor-C (VEGF-C), monocyte chemotactic protein-1 (MCP-1), interleukin-1β (IL-1β), IL-4, IL-6, IL-7, IL-10, IL-17, interferon-γ (IFN-γ), and transforming growth factor-β (TGF-β) (*6, 8, 15-20*). Therefore, we determined their concentrations in the CSF and serum of each group of mice. In the CSF, both nasal-group and i.c./nasal-group had a significant increase in the level of VEGF-C, MCP-1, IFN-γ and TGF-β compared with con-group (Fig.6). The i.c./nasal-group possessed a significant larger extent of increase than the nasal-group (Fig.6). The other tested cytokines in the CSF were of no significant differences among groups or undetectable in all groups. In serum, both nasal-group and i.c./nasal-group had a significant increase in the level of MCP-1, IFN-γ, TGF-β, IL-4 and IL-10 compared with con-group (Fig.7). The i.c./nasal-group possessed a significant larger extent of increase than the nasal-group (Fig.7). The other tested cytokines in serum were of no significant differences among groups or undetectable in all groups.

**Fig.6.**
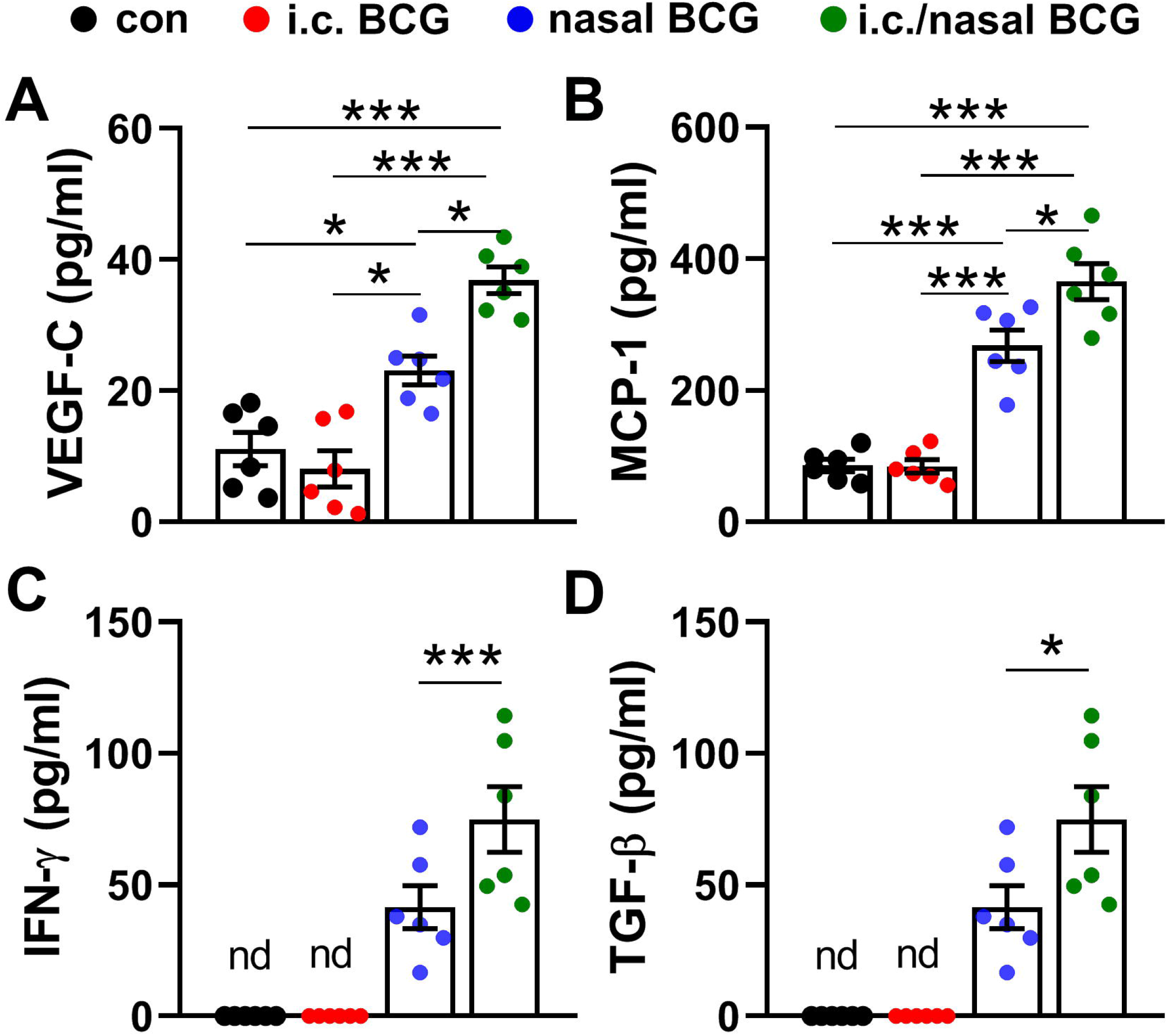
Elevated cytokine levels in CSF in nasal-group and i.c./nasal-group at P21. (A-B) Bars represent cytokines levels. *n* = 6 mice/group. **: *p* < 0.01; ***: *p* < 0.001. Randomly completely block design ANOVA followed by Bonferroni’s *post hoc* test. (C-D) Representative bars represent cytokines levels. *n* = 6 mice/group. nd indicates undetectable levels of one cytokine in certain group, such as con-group; ***: *p* < 0.001. Paired-samples *t* test between nasal-group and i.c./nasal-group. Data are presented as the mean ± SEM.

**Fig.7.**
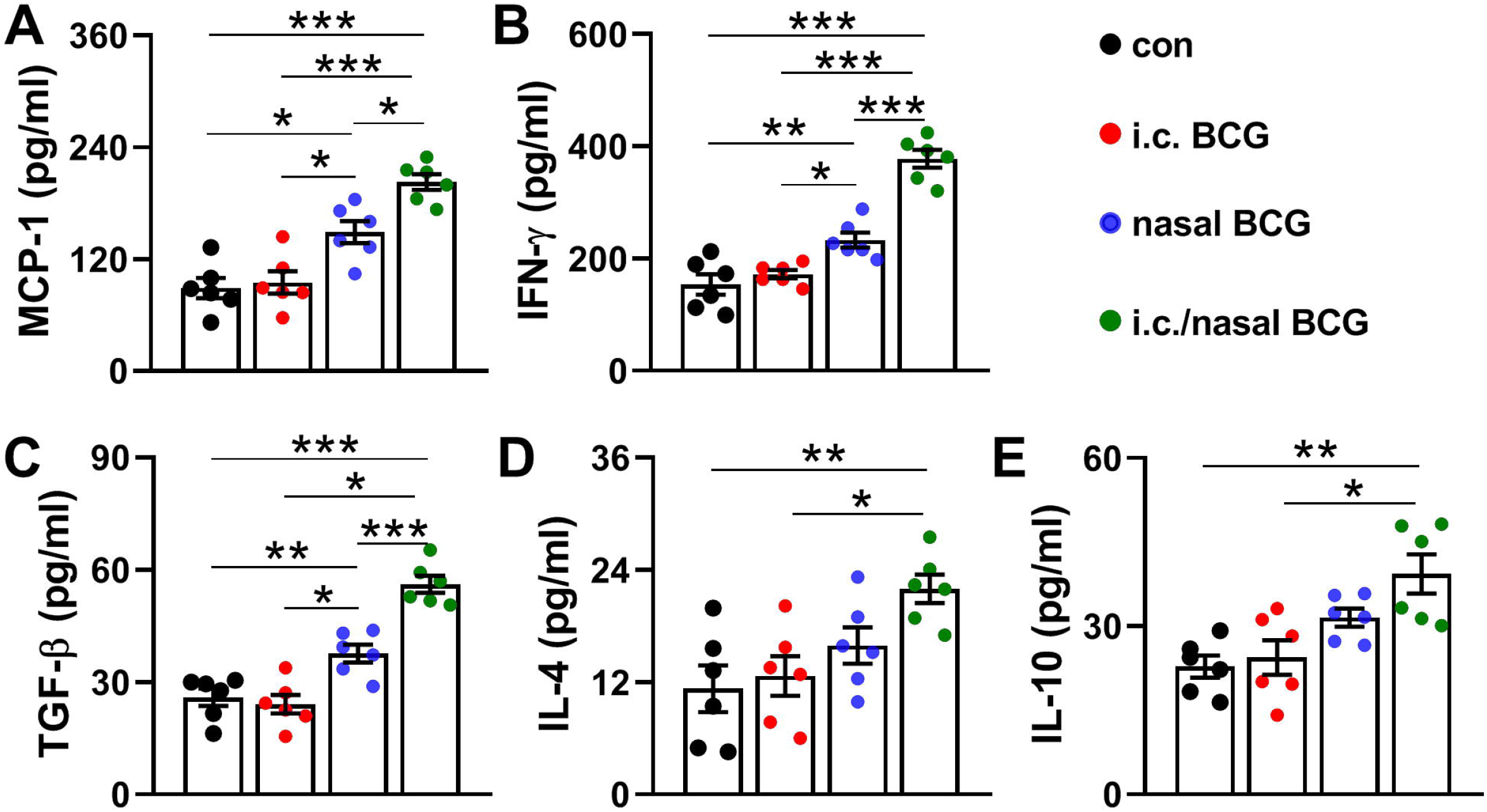
Elevated cytokine levels in serum in nasal-group and i.c./nasal-group at P21. (A-B) Bars represent cytokines levels. *n* = 6 mice/group. *: *p* < 0.05; **: *p* < 0.01; ***: *p* < 0.001. Randomly completely block design ANOVA followed by Bonferroni’s *post hoc* test. Data are presented as the mean ± SEM.

Besides, the level of VEGFR-3, the receptor for VEGF-C and mainly expressed by lymphatics, was also detected in dura mater because of its vital role in lymphangiogenesis (*21, 22*). WB analysis showed both nasal-group and i.c./nasal-group had a significant increase in VEGFR-3 expression (Fig. 8). In both indexes, the i.c./nasal-group possessed a significant larger extent of increase than the nasal-group (Fig.8).

**Fig.8.**
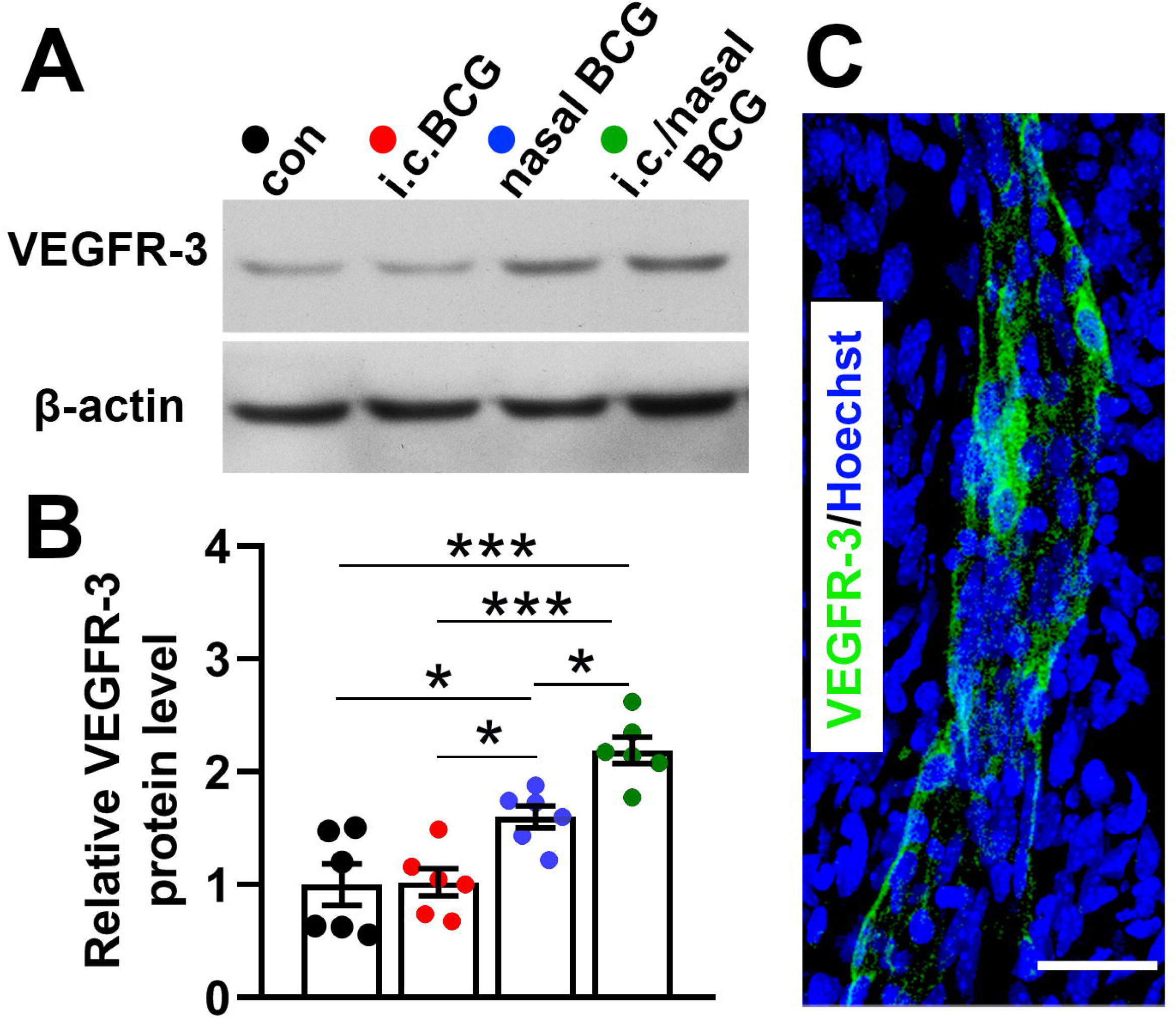
Increased VEGFR-3 protein level in dura mater of the mice in nasal-group and i.c./nasal-group. (A) Representative western blots showing VEGFR-3 protein level in dura mater of each group. (B) Quantitative data showing alteration in the relative levels of the VEGFR-3 protein in the dura mater. *n* = 6 mice/group. *: *p* < 0.05; **: *p* < 0.01; ***: *p* < 0.001. Randomly completely block design ANOVA followed by Bonferroni’s *post hoc* test. Data are presented as the mean ± SEM. (C) Representative graphs show VEGFR-3 expression in dLVs. Scale bar, 50 μm.

### Macrophage identified as a mediator of BCG-induced promotion in dural lymphatics development

VEGF-C, mainly secreted by macrophages, is the most important cytokine for promoting lymphangiogenesis (*16, 17, 23*). MCP-1 is the most potential chemokine for recruting macrophages (*24*). Therefore, the increased levels of them in the CSF in nasal-group and i.c./nasal-group may explain the larger number of macrophages counting in nasal-group and i.c./nasal-group as shown in Fig.2H, Fig.3 and Fig. S1.

To further confirm the mediating role of macrophages in BCG-induced promotion in dural lymphatics development, each group of mice were intraperitoneally injected with a macrophage-depleting agent clodronate liposomes (110 mg/kg, once every three days). At P28, dura mater were stained for Lyve-1^+^ vessels. The mean Lyve-1^+^ vessels area in SSS zone in CL/con-group (Fig.S2D) is less than that of con-group at P21 (Fig.2A, Fig. S1A). Moreover, there is no longer any significant difference among four groups in either Lyve-1^+^ vessels area in SSS zone or in the level of VEGF-C in CSF after macrophage-depletion (Fig.S2). What’s more, rare Lyve-1^+^ cells out of Lyve-1^+^ vessels could be observed in dura mater when macrophages were depleted.

## Discussion

This study for the first time revealed that repeated neonatal BCG vaccination, either combined with or without i.c. injection of BCG, accelerated dural lymphatics development in mice. Furthermore, macrophages were identified as the mediator of BCG-induced dural lymphatics development acceleration.

A notable phenomenon found in this study is that a single dose of BCG i.c. injection resulted in no significant influence in dural lymphatics development at P21.

This may be because of the lack of boosting immunization. The possible role of boosting immunization was also supported by the fact that the i.c./nasal-group consistently possessed a significant larger extent of alteration than the nasal-group in nearly all indexes. The present study aimed mainly to explore the potential influence of the immune activation pattern ‘neonatal i.c. injection/nasal immunization of *mycobacterium tuberculosis’* that happens to people worldwide. Therefore, boosting immunization by repeating i.c. injection of BCG was not attempted in this study.

BCG immunization could not only induce Th1 cytokines production such as IFN-γ (*25*) but also activate macrophages (*26*). This was confirmed by the results of cytokines analysis in this study. Macrophages is the major source of VEGF-C protein (*16*). In the nasal-group and i.c./nasal-group, the elevated VEGF-C levels in the CSF (Fig.6A) may be explained by the increased numbers of macrophages in the dura mater. In turn, the larger number of recruited macrophages in the dura mater may be explained by the higher CSF level of MCP-1 that is a very potential chemokine for monocyte/macrophages (*27*).

It is reported that BCG could induce systemic release of MCP-1 (*28*). However, although the base level of MCP-1 both in serum and CSF indicated by the data from con-group (Fig.6B and Fig.7A), the CSF MCP-1 level was about twice of its serum level in nasal-group and i.c./nasal-group. This suggests that the MCP-1 in the CSF may not derive from circulating blood, but rather produced by local cells. Activated microglia was verified to serve as a possible producer of MCP-1 (*29*). Neonatal BCG immunization has been shown to activate microglia (*25, 30*). This deduction provides with an interest cue for future research to investigate the exactly mechanism under the role of BCG in increasing the CSF level of MCP-1.

According to the previous report, the frontiers of the growth point of dural lymphatics reach the scalp of skull such as the lateral parts of TS area at about two weeks old (*5*). Given that the nasal BCG vaccination lasted for two weeks or more, the dura mater of TS and SSS area, not skull base, was chosen for dural lymphatics analysis. Moreover, the complex anatomical structure of the base of skull and the tight attaching of dura mater there would make it difficult to analyze the lymphatics precisely. It is similar with the part of dura mater overlapping attached to the frontal bone overlying the olfactory bulb. In addition, the part of dura mater attached to the scalp of skull is easy to be isolated from bone.

The dural lymphatics begin to cover the whole length area along the SSS but are often discontinuous even in adult mice as reported previously (*5*). These vessels in mice at P21 were seen in about 1/4 to 1/3 of the whole SSS length in con-group mice (Fig. 2A). At P21, mere nasal BCG vaccination mildly increased the dural lymphatics whereas mere neonatal i.c. BCG injection brought no significant effects. Both the nasal-group and i.c./nasal-group showed an advanced development of the dural lymphatics at P21, with their dural lymphatics distribution reaching the level comparable with those in normal adult mice. This advanced development during the critical period for CNS development is very likely to allow an advanced and more effective drainage of CSF and thereby to produce an affect on brain development.

In natural condition, the microbiota to which human beings nasally expose are certainly different from nasal BCG vaccination used in this study both in kind and in dosage. Admittedly, it is not proper to estimate the status of dural lymphatics development in human beings straightly according to the findings in this study especially given that it is not clear whether the dural lymphatics development in perinatal period differs in human beings from that in mice. What is vital and novel in this study is that nasal BCG vaccination does promote dural lymphatics development. Even if this promotion means only a matter of an advanced development or an earlier coming of matured dural lymphatics, rather than a long-lasting elevation of lymphatics structure and function, it is still important for brain development since it occurs during the the early life period when is critical for CNS development programming. In other words, even a transient promoted/advanced dural lymphatics development is likely to exert a long-lasting effects on brain development and function. In addition, the protocol used in this work to immunize mice may serve as an animal model for experiments studying dural lymphatics-related issues.

## Authors’ contributions

Study design: Junhua Yang and Zhibin Yao. Performing the experiments: Junhua Yang (primarily morphological observations and ELISA), Lifang Yuan (primarily animal treatments and grouping, morphological observations and ELISA), Linyang Song (primarily multiplex assays and western blotting). Data analyses (blinded to groups): Junhua Yang. Writing the paper: Junhua Yang. Funding acquisition: Junhua Yang, Zejie Zuo and Zhibin Yao. Fangfang Qi and Jie Xu participated in discussions, assisted with data analysis, technical instructions and manuscript editing. Conceptualization, Supervision All authors read, commented and approved the final manuscript.

## Conflict of interest

The authors declare that they have no conflict of interest.

## Acknowledgments

This work was supported by grants from the starting fund for high-level talent introduction into Guangdong Pharmaceutical University (No. 51355093) to Junhua Yang, the Special Foundation of Education Department of Guangdong Province, the Medical Scientific Research Foundation of Guangdong Province, China (2013-159) to Zhibin Yao and the National Natural Science Foundation of China (Nos. 81801058) to Zejie Zuo. The funding agencies had no role in study design and performance; in the collection, analysis and interpretation of data; in the writing of the report; or in the decision to submit the paper for publication. We thank Technician Yuanjun Guan (Zhongshan School of Medicine, SYSU) for her kind assistance with experimental techniques, particularly the confocal imaging and analyses.

## Notes

### Competing Interest Statement

The authors have declared no competing interest.

### Summary of Updates

More detailed information of the affiliations of all authors has been revised in this submitted version. No change was made in the content of all the manuscript files.

