## Supplementary figures and legends for "Nasal BCG Exposure Accelerates Dural Lymphatics Development via Macrophages’ Role in Newborn Mice"

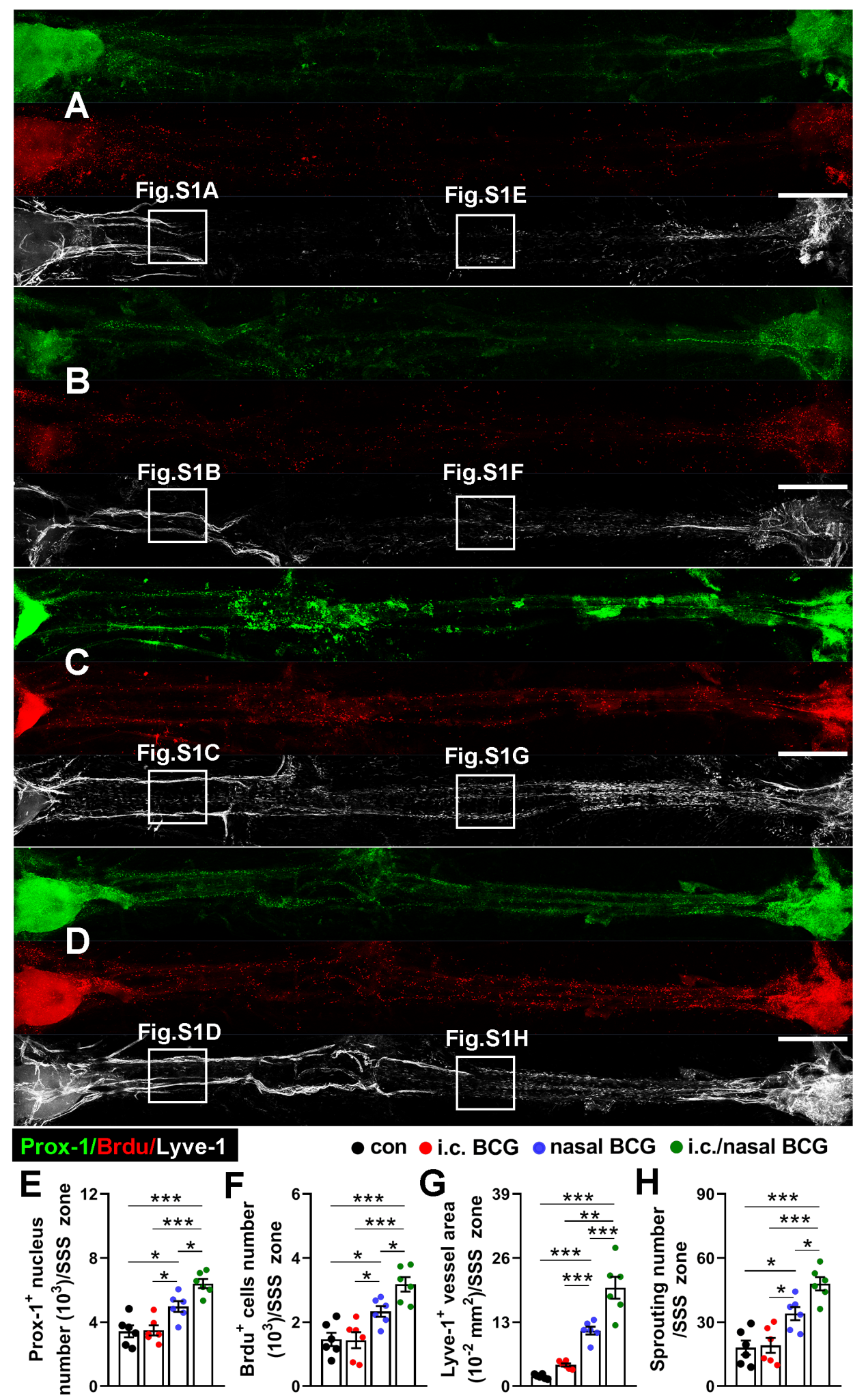

**Fig.S1 Elevated real time lymphangiogenesis level in SSS zone in nasal-group and i.c./nasal-group at P19** (A-D) Representative graphs show the differences in dLVs distribution (white) as well as the BrdU^+^ (red) and Prox-1^+^ (green) cells intensity within the corresponding SSS areas of the dura in each group of mice. Scale bar, 400 μm. (E-H) Bars represent Prox-1^+^ nucleus numbers, BrdU^+^ nucleus numbers, Lyve-1^+^ vessels area% and numbers of sprouting from Lyve-1^+^ vessels. *n* = 6 mice/group. *: *p* < 0.05; **: *p* < 0.01; ***: *p* < 0.001. Randomly completely block design ANOVA followed by Bonferroni’s *post hoc* test. Data are presented as the mean ± SEM.

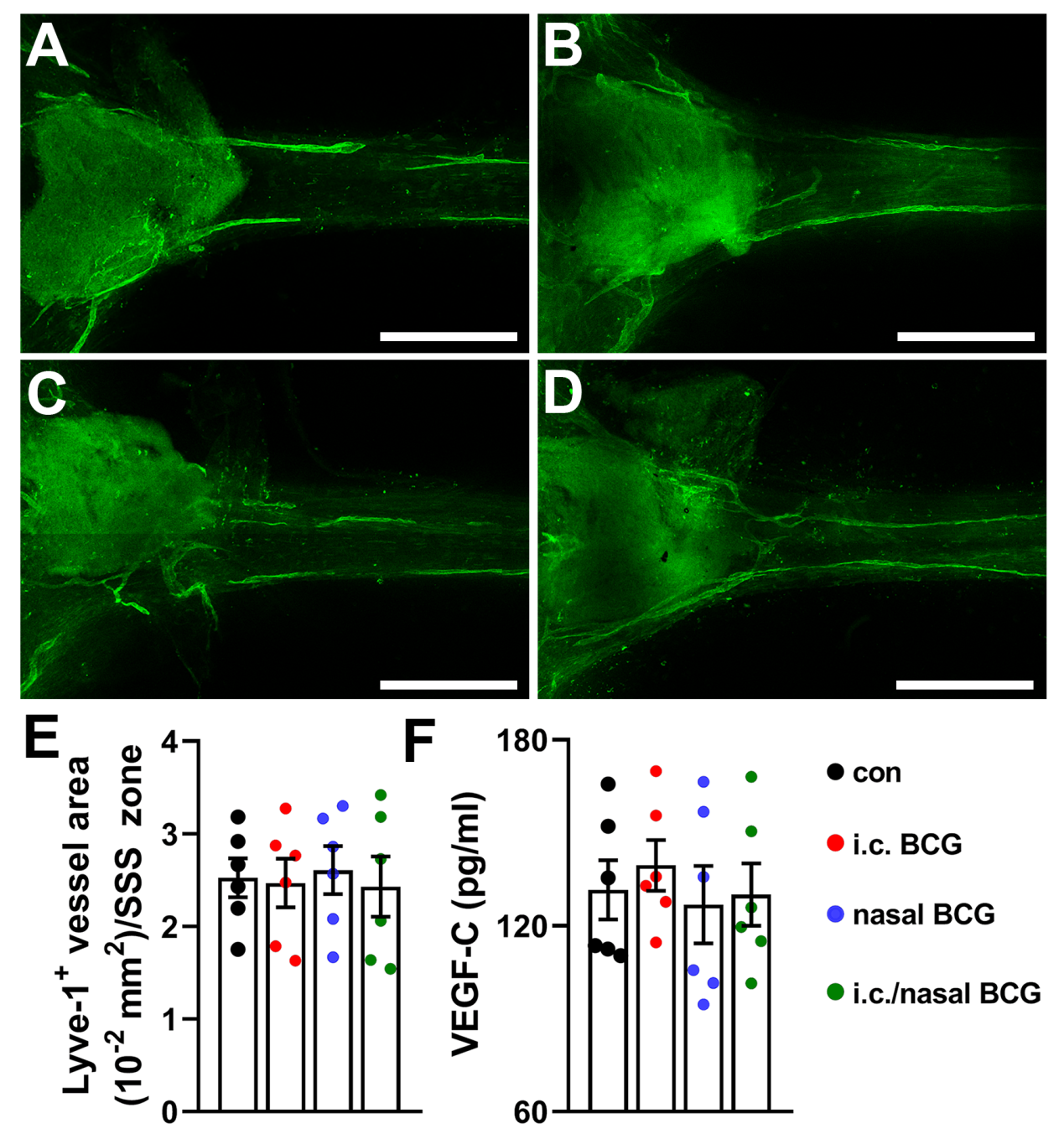

**Fig.S2 Macrophage identified as a mediator of BCG-induced promotion in dural lymphatics development.** (A-D) Representative graphs show Lyve-1^+^ vessels (dLVs) distribution within the corresponding SSS areas of the dura in each group of mice. Scale bar, 400 μm. (E) Bars represent Lyve-1^+^ vessels area quantification in each group of mice. (F) Bars represent CSF VEGF-C quantification in each group of mice. *n* = 6 mice/group. Randomly completely block design ANOVA followed by Bonferroni’s *post hoc* test. Data are presented as the mean ± SEM.
